# Imaging cytoplasmic lipid droplets *in vivo* with fluorescent perilipin 2 and perilipin 3 knockin zebrafish

**DOI:** 10.1101/2021.01.10.426109

**Authors:** Meredith H. Wilson, Stephen C. Ekker, Steven A. Farber

## Abstract

Cytoplasmic lipid droplets are highly dynamic storage organelles; their rapid synthesis, expansion, and degradation, as well as their varied interactions with other organelles allow cells to maintain lipid homeostasis. While the molecular details of lipid droplet dynamics are currently a very active area of investigation, this work has been primarily performed in cultured cells and *in vitro* systems. By taking advantage of the powerful transgenic and *in vivo* imaging opportunities afforded by the zebrafish model system, we have built a suite of tools to allow lipid droplets to be studied in real-time from the subcellular to the whole organism level. Fluorescently-tagging the lipid droplet associated proteins, perilipin 2 and perilipin 3, in the endogenous loci, permits visualization of lipid droplets in the intestine, liver, lateral line and adipose tissue. Using these transgenic lines we have found that perilipin 3 is rapidly loaded on intestinal lipid droplets following a high fat meal and then largely replaced by perilipin 2 a few hours later. These powerful new tools will facilitate studies on the role of lipid droplets in different tissues and under different genetic and physiological manipulations.

## Introduction

Cytoplasmic lipid droplets are cellular organelles composed of a core of neutral lipids surrounded by a monolayer of phospholipids and coated with a variety of proteins. While initially believed to be passive storage depots for lipids, it is now appreciated that lipid droplets are dynamic organelles with roles in cellular lipid homeostasis, protection from lipotoxicity and ER stress, viral and parasitic infection, and in host defense (Bosch et al., 2020; Cloherty et al., 2020; Coleman, 2020; Farese and Walther, 2009; Olzmann and Carvalho, 2019; Roberts and Olzmann, 2020).

Lipid droplets are typically coated by one or more perilipins, an evolutionarily related protein family defined by two conserved protein motifs, the N-terminal ~100 amino acid hydrophobic PAT domain followed by a repeating 11-mer helical motif of varying length (Kimmel and Sztalryd, 2016). Perilipins (PLINs) are recruited to the lipid droplet surface directly from the cytosol, mediated at least in part by the 11-mer repeat regions which fold into amphipathic helices (Kimmel and Sztalryd, 2016; Rowe et al., 2016). Perilipins act to regulate lipid storage by through their role in preventing or modulating access of lipid droplets to lipases and lipophagy (Sztalryd and Brasaemle, 2017).

Perilipins have been found in species ranging from *Dictyostelium* to mammals, with more divergent variants in fungi and *Caenorhabditis* (Bickel et al., 2009; Gao et al., 2017; Kimmel and Sztalryd, 2016; Sztalryd and Brasaemle, 2017). The human genome contains five perilipin genes, now designated PLIN1 – 5. Perilipins 2 and 3 are expressed ubiquitiously (Brasaemle et al., 1997; Diaz and Pfeffer, 1998; Heid et al., 1998; Wolins et al., 2001), whereas perilipin 1 is predominantly expressed in white and brown adipocytes (Greenberg et al., 1991; Lu et al., 2001). PLIN4 is expressed in adipocytes, brain, heart and skeletal muscle, and PLIN5 is found in fatty acid oxidizing tissues such as heart, brown adipose tissue, and skeletal muscle, as well as in the liver (Dalen et al., 2007; Wolins et al., 2006; Yamaguchi et al., 2006). The genomes of rayfin fish, including zebrafish, have orthologs of human PLIN1, PLIN2 and PLIN3 in addition to a unique PLIN variant, perilipin 6 which targets the surface of pigment-containing carotenoid droplets in skin xanthophores (Granneman et al., 2017).

While fluorescently-tagged perilipin reporter proteins are used extensively in cell culture to visualize lipid droplets (for example (Chung et al., 2019; Granneman et al., 2017; Kaushik and Cuervo, 2015; Miura et al., 2002; Schulze et al., 2020; Targett-Adams et al., 2003)), lipid droplets *in vivo* have been historically studied in fixed tissues using immunohistochemistry (Frank et al., 2015; Lee et al., 2009), staining with lipid dyes (Oil red O, Sudan Black, LipidTox), or by electron microscopy (Chughtai et al., 2015; Marza et al., 2005; Zhang et al., 2010). Lipid droplets can also be labeled in live organisms with fluorescent lipophilic dyes such as BODIPY (Mather et al., 2019) & Nile red (Minchin and Rawls, 2017b), fed with fluorescently-tagged fatty acids (BODIPY & TopFluor) which are synthesized into stored fluorescent triglycerides or cholesterol esters (Ashrafi et al., 2003; Carten et al., 2011; Furlong et al., 1995; Quinlivan et al., 2017), or imaged in the absence of any label using CARS or SRS microscopy (Chien et al., 2012; Chughtai et al., 2015; Wang et al., 2011). However, expression of fluorescently-tagged lipid droplet associated proteins *in vivo* has been primarily limited to yeast (Gao et al., 2017), *Drosophila* (Bi et al., 2012; Gronke et al., 2005) and *C. elegans* (Chughtai et al., 2015; Xie et al., 2019; Zhang et al., 2010), although a transgenic zebrafish *plin2-tdtomato* line was very recently described (Lumaquin et al., 2020).

Here, we report the generation and validation of zebrafish perilipin reporter lines, including *Fus*(*EGFP-plin2*) and *Fus*(*plin3-RFP*), in which we inserted fluorescent reporters in-frame with the coding sequence at the genomic loci. These reporter lines faithfully recapitulate the endogenous expression of *plin2* and *plin3* in larval zebrafish, allowing for *in vivo* imaging of lipid droplet dynamics in live animals at the subcellular, tissue, organ and whole larvae level. Using these lines, we describe the ordered recruitment of plin3 and then plin2 to lipid droplets in intestinal enterocytes following the consumption of a high fat meal, reveal a delay in hepatic expression of plin2 and plin3 during development and identify a population of plin2-positive lipid droplets adjacent to neuromasts in the posterior lateral line.

## Results and Discussion

### *Perilipin 2* and *perilipin 3* are expressed in the intestine of larval zebrafish

To determine the tissue localization of *plin2* and *plin3* mRNA expression in larval zebrafish, we performed whole mount *in situ* hybridization. Our data indicate that *plin2* is not expressed in any tissues of unfed larvae at 6 days post fertilization (dpf) (Figure 1A,1B). However, following a high-fat meal, *plin2* mRNA expression is strongly induced in the intestine, consistent with findings in mice (Lee et al., 2009) and with our previous RNAseq and qRT-PCR data (Zeituni et al., 2016). *plin3* mRNA is present in the intestines of both unfed and fed larvae, and the signal in the intestine is stronger in high-fat fed larvae (Figure 1A & 1B), again consistent with previous data in fish and mice (Lee et al., 2009; Zeituni et al., 2016). Surprisingly, neither *plin2* nor *plin3* mRNA expression was noted in the liver or in other tissues of unfed or high-fat fed zebrafish larvae at 6 dpf (Figure 1A & 1B) as was expected from studies in mouse and human tissues (Brasaemle et al., 1997; Heid et al., 1998; Than et al., 1998; Wolins et al., 2006).

**Figure 1:**
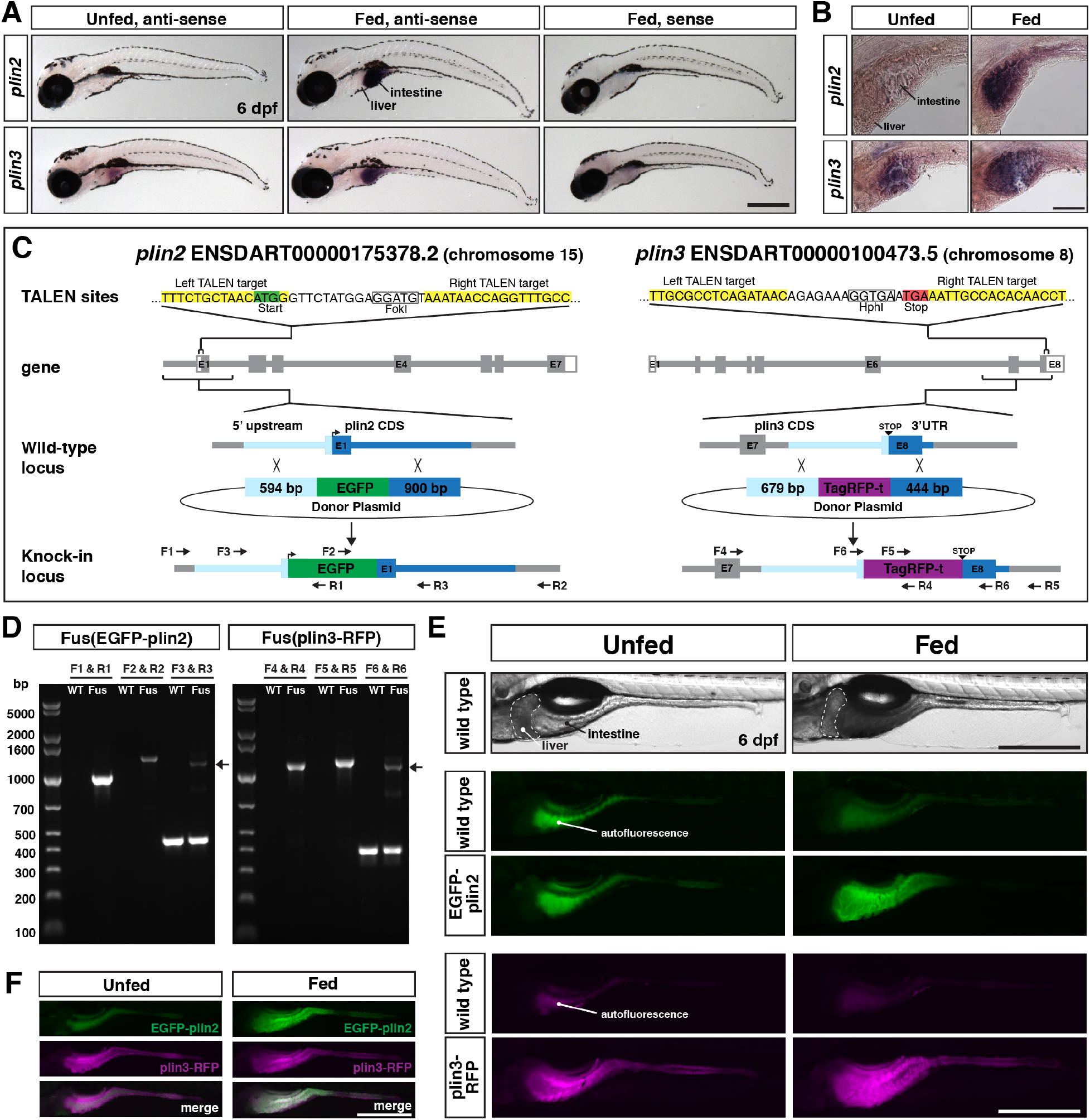
Generation of in-frame fluorescent reporters in the endogenous *plin2* and *plin3* loci. (A,B) Representative images of whole mount *in situ* hybridization with probes against zebrafish *plin2* (*ENSDARG00000042332*) and zgc:77486 (*plin3/4/5*) (*ENSDARG00000013711*) at 6 dpf either unfed or following feeding with a high fat meal for 90 min. ISH was performed 3 times for each gene with n = 10 larvae per probe per experiment; scale = 500 μm (A), scale = 100 μm (B). *Plin2* is expressed in the intestine only following a high-fat meal whereas *plin3* is expressed in the intestine in unfed fish and has stronger expression following a high fat meal. (C) Overview of the location and strategy used for TALEN-mediated genome editing. EGFP was fused in-frame at the N-terminus of *plin2*. TALEN targets in *plin2* are located in exon 1 of the *plin2-203 ENSDART00000175378.2* transcript and flank a FokI restriction site, loss of which was used to confirm cutting activity. A donor plasmid with the coding sequence for EGFP and *plin2* homology arms was co-injected with TALEN mRNA into 1-cell stage embryos to be used as a template for homology directed repair. mTag-RFP-t was fused in frame at the C-terminus of *plin3*. TALEN targets were located in exon 8 of the *plin3 ENSDART00000100473.5* transcript and flank the termination codon and a HphI restriction site, loss of which was used to confirm cutting activity. A donor plasmid with the coding sequence for mTagRFP-t and *plin3* homology arms was co-injected with TALEN mRNA into 1-cell stage embryos to be used as a template for homology directed repair. (D) Following identification of fluorescent embryos in the F1 generation, RT-PCR and sequencing of genomic DNA using the primers noted on the knock-in loci depicted in (C) were used to confirm successful in-frame integration of the fluorescent tags. The size of the amplicons expected for correct integration were as follows: F1-R1 1033bp, F2-R2 1340bp, F3-R3 440bp for WT & 1224bp for *Fus*(*EGFP-plin2*) fusion, F4-R4 1218bp, F5-R5 1274bp, F6-R6 401bp for WT & 1187 for *Fus*(*plin3-RFP*). Arrows indicate the larger amplicon in heterozygous fish carrying the fusion alleles. (E) Imaging in live larvae (6 dpf) reveals expression of EGFP-plin2 only in the intestine of larvae fed a high-fat meal (7 h post-start of 2 h meal) and plin3-RFP is expressed in the intestine of both unfed and fed larvae (4.5 h post-start of 2 h meal, larvae are heterozygous for the fusion proteins; the lumen of the intestine has strong autofluorescence in wild-type and transgenic fish; see Figure 1 – figure supplement 2 for images of whole fish). Scale = 500 μm. (F) Examples of larvae expressing both EGFP-plin2 and plin3-RFP (7 h post start of meal). Scale = 500 μm.

### Generation of knock-in/fusion lines to study lipid droplets *in vivo*

To study how plin2 and plin3 regulate lipid droplet dynamics *in vivo*, we generated fluorescent *plin2* and *plin3* zebrafish reporter lines (Figure 1C). We specifically wanted to explore the precise timing of plin2 and plin3 association with lipid droplets immediately following a high fat meal. We also recognized that overexpression of plin proteins can alter lipid droplet dynamics by altering the rate of lipolysis and lipophagy, which can result in altered levels of cytoplasmic lipid and lipoprotein secretion (Bell et al., 2008; Bosma et al., 2012; Fukushima et al., 2005; Listenberger et al., 2007; Magnusson et al., 2006; McIntosh et al., 2012; Tsai et al., 2017). Therefore, we chose to tag the endogenous proteins by engineering knock-in alleles. We used TALENs to introduce a double strand break near the start codon in exon 1 in *plin2* (*ENSDART00000175378.2*) or adjacent to the termination codon in the last exon of *plin3* (*ENSDART00000100473.5*). TALEN mRNA was injected into 1-cell stage embryos, together with donor constructs including either *EGFP* for plin2 or *tagRFP-t* for *plin3*, flanked by the noted homology arms to direct homology directed repair (Figure 1C). The left homology arm for *plin2* included the 54-bp variable sequence we discovered upstream of exon 1, that may contain a regulatory element for control of *plin2* expression (Figure 1 – figure supplement 1). From the injected F0 adult fish, we identified a single founder for each knock-in allele by out-crossing and screening progeny for fluorescence in the intestine at 6 dpf either following a meal (*Fus*(*EGFP-plin2*)) or prior to feeding (*Fus*(*plin3-RFP*)). Correct integration of the fluorescent constructs was confirmed by PCR on genomic DNA of individual larvae, followed by sequencing (Figure 1D).

Consistent with our whole mount *in situ* hybridization data, we observe EGFP-plin2 only in the intestine of larvae fed a high-fat meal, whereas plin3-RFP is detected in the intestine of both unfed and fed larvae (Figure 1E). No fluorescence is noted in the liver or in other tissues at 6 dpf, regardless of feeding status (for images of whole fish see Figure 1 – figure supplement 2). Fish carrying the knock-in alleles can be in-crossed and resulting progeny express RFP-plin3 in the intestine prior to feeding and both transgenes are expressed subsequent to consuming a high-fat meal (Figure 1F). Thus, these knock-in alleles faithfully recapitulate the endogenous tissue mRNA expression patterns of *plin2* and *plin3* as revealed by in situ hybridization of fixed larval zebrafish (Fig 1A).

### EGFP-plin2 and plin3-RFP decorate lipid droplets in intestinal enterocytes

To confirm that EGFP-plin2 and plin3-RFP decorate the surface of lipid droplets in the intestine, we performed confocal imaging in live larvae following a high-fat meal. *Fus(EGFP-plin2)/+* larvae were fed liposomes containing the fluorescent fatty acid analogue BODIPY 558/568-C12. This fatty acid analogue can be incorporated into both phospholipid and triglycerides for storage in lipid droplets in larval zebrafish (Quinlivan et al., 2017). As expected, EGFP-plin2 decorates the surface of BODIPY 558/568-C12-positive lipid droplets in the intestine, which is visualized as rings of EGFP fluorescence in single confocal z-slices (Figure 2A). Similarly, plin3-RFP localizes to the surface of intestinal lipid droplets labeled with the green fluorescent BODIPY FL-C12 fatty acid analogue (Figure 2A). In larvae heterozygous for both *Fus*(*EGFP-plin2*) and *Fus*(*plin3-RFP*), we found that lipid droplets can be labeled by both plin3-RFP and EGFP-plin2 proteins (Figure 2B). While resolving cell membranes in the anterior intestine of larvae is difficult with bright-field or differential interference contrast microscopy due to the three-dimensional nature of the intestinal folds and small cell size, these perilipin reporter lines could be crossed to fish carrying transgenic markers of the cell membranes (Alvers et al., 2014) to better elucidate to localization of lipid droplets within individual enterocytes if desired.

**Figure 2:**
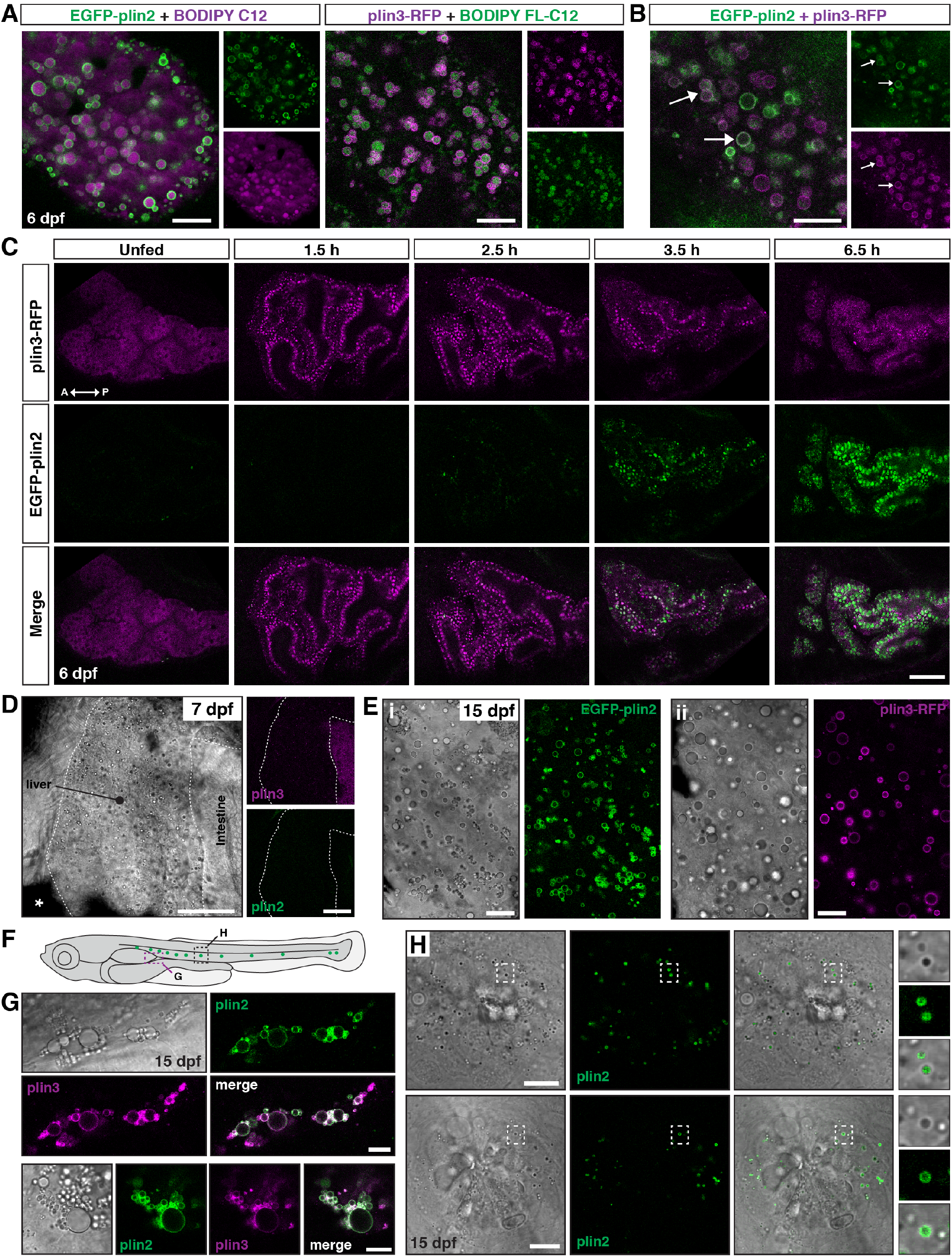
EGFP-plin2 and plin3-RFP decorate lipid droplets in the intestine, liver, adipocytes and in cells surrounding neuromasts. (A) EGFP-plin2 (green) and plin3-RFP (magenta) label the lipid droplet surface in the intestine of 6 dpf larvae fed with a high-fat meal containing either BODIPY 558/568 C12 (magenta) or BODIPY FL-C12 (green) to label the stored lipids. Note the 558/568 C12 is not fully incorporated into stored lipid and is also found diffuse in the cytoplasm. Scale = 10 μm. (B) EGFP-plin2 and plin3-RFP can decorate the same lipid droplets in the intestine. Arrows denote examples of dual-labeled droplets, scale = 10 μm. (C) Lateral views of the anterior intestine in unfed larvae and in larvae at different time-points following the start of feeding with a high-fat meal for 1 h. Fish were heterozygous for both *Fus*(*plin3-RFP*) and *Fus*(*EGFP-plin2*). Images are representative of 3 independent experiments (15-25 fish per experiment); data presented are from one experiment. Scale = 50 μm. (D) Lateral view of the liver in a 7 dpf larvae heterozygous for both *Fus*(*plin3-RFP*) and *Fus*(*EGFP-plin2*). Scale = 50 μm. (E) Liver micrographs from 15 dpf larval zebrafish fed a diet of Gemma + 4% cholesterol for 10 days. Lipid droplets in hepatocytes can be labeled with EGFP-plin2 (i) and with plin3-RFP (ii). Scale = 20 μm. (F) Cartoon of 15 dpf larval zebrafish showing the general location of images in panels G and H. (G) EGFP-plin2 and plin3-RFP can both decorate lipid droplets in adipocytes. Fish are heterozygous for both transgenes and were fed Gemma + 4% cholesterol for 10 days. Scale bars = 10 μm. (H) Examples of lipid droplets around neuromasts in fish heterozygous for *Fus*(*EGFP-plin2*) at 15 dpf. Panels on right show droplets in the boxed regions to the left. Scale bars = 10 μm.

### Investigating the temporal dynamics of EGFP-plin2 and plin3-RFP in live intestinal enterocytes

The localization of perilipin 2 and perilipin 3 on lipid droplets in enterocytes in response to high fat remains poorly understood. Prior studies in mice have suggested that PLIN3 is located on lipid droplets following an acute high-fat meal, but not following chronic high-fat feeding (D’Aquila et al., 2015; Lee et al., 2009). In contrast, despite upregulation of PLIN2 protein in the intestine following an acute feed, PLIN2 was only present on lipid droplets in enterocytes following chronic high-fat feeding (D’Aquila et al., 2015; Lee et al., 2009). However, these findings were based on single time-points and no studies have imaged perilipins in the intestine at the level now possible with these transgenic zebrafish lines.

Plin3 is expressed throughout the cytoplasm in cells, including enterocytes, in the absence of lipid droplets and binds to nascent lipid droplets as they emerge from the endoplasmic reticulum (Bulankina et al., 2009; Chung et al., 2019; Lee et al., 2009; Skinner et al., 2009; Wolins et al., 2005). In contrast, plin2 is only stable when bound to lipid droplets, and is quickly ubiquitinated and degraded in the absence of droplets (Xu et al., 2005). In keeping with these data, in unfed larval zebrafish, in the absence of lipid droplets, plin3-RFP is distributed throughout the cytoplasm of the intestinal enterocytes (Figure 2C, Unfed). After consuming a high-fat meal, the plin3-RFP pattern changes considerably, there is less cytoplasmic signal and bright puncta are present, which correspond to lipid droplets of various sizes (1.5 h). However, weak EGFP-plin2 signal, exclusively on lipid droplets, only emerges ~2-3 hr after the start of a meal, which is consistent with the need to both transcribe (Figure 1A & (Zeituni et al., 2016)) and translate the protein after the start of the meal. As time continues, the EGFP-plin2 signal on droplets increases strongly and the plin3-RFP fluorescence becomes predominantly cytoplasmic again by 6-7 h.

Because plin3-RFP is already present in the cytoplasm before the emergence of lipid droplets, the timing of the appearance of this transgene on lipid droplets is likely similar to the unlabeled plin3 allele. However, while we suspect that the additional time necessary to translate and fold the EGFP tag (~30-60 min, (Balleza et al., 2018; Heim et al., 1995)) likely delays the appearance of EGFP-plin2 on lipid droplets compared to unlabeled plin2, the visible detection of EGFP-plin2 starting at ~2-3 h is still consistent with the peak in *plin2* mRNA expression between 1-3 h following the start of a high-fat meal (Zeituni et al., 2016). It is also unclear whether the fluorescent tags alter the stability or removal of the perilipins from the lipid droplets. Despite these caveats, these reporters indicate a clear progression from plin3 to plin2 on intestinal lipid droplets in zebrafish. The shift from plin3 to plin2 over time is consistent with displacement of PLIN3 by PLIN2 as lipid droplets grow in size in cultured 3T3-L1 adipocytes (Wolins et al., 2005). However, it is unclear whether this ordered recruitment in the intestine is specific to fish, or whether it would also be observed in mammals if more frequent sampling was performed following an acute high-fat meal. The physiological significance of this shift in perilipin lipid droplet association in enterocytes remains to be determined, though we hypothesize that it may aid in the regulation of chylomicron production and post-prandial plasma lipid levels.

### Plin2 and plin3 decorate hepatic lipid droplets in older larvae

Brightfield imaging of livers in larvae at 6-7 dpf suggest they contain lipid droplets (Figure 2D) and we have shown previously that these droplets store lipids synthesized with BODIPY fatty acid analogues (Carten et al., 2011; Quinlivan et al., 2017). However, confocal imaging in unfed and fed fish indicates that these droplets are only very rarely labeled with EGFP-plin2 or plin3-RFP (Figure 2D). In contrast, hepatic lipid droplets in older larvae can be labeled with EGFP-plin2 and plin3-RFP (Figure 2E). While labeled lipid droplets are present in fish fed our standard Gemma diet, when this diet was supplemented with 4% cholesterol, the hepatic lipid droplets tend to be more abundant and are more often decorated by perilipins 2 and 3 (Figure 2E). Studying the differential expression of perilipins in hepatocyte of young vs. older larvae using these transgenic lines provide an opportunity to yield novel insights into the transcriptional regulation of perilipins and how the perilipins influence hepatic lipid storage and mobilization *in vivo*.

### EGFP-plin2 and plin3-RFP decorate adipocyte lipid droplets

Using fluorescent lipophilic dyes, Minchin & Rawls have described 34 distinct regions of adipose tissue in zebrafish, including 5 visceral and 22 subcutaneous adipose tissue depots (Minchin and Rawls, 2017a; Minchin and Rawls, 2017b). One of the earliest depots to develop is the abdominal visceral adipose tissue (AVAT), which appears at the posterior aspect of the swim bladder at ~5 mm standard length (Minchin and Rawls, 2017b). Confocal imaging in fish carrying the *plin* knock-in alleles indicates that the lipid droplets in the adipocytes of the AVAT tissue are labeled with both EGFP-plin2 and plin3-RFP (Figure 2F & 2G). Thus, we expect that the perilipin knock-in alleles will be a valuable tool to assist in studies of adipocyte lipid droplet dynamics *in vivo* during development and in pathological conditions.

### EGFP-plin2 indicates the presence of lipid droplets around neuromasts

Unexpectedly, when imaging larvae at 15 dpf, we consistently noted small EGFP-plin2-positive lipid droplets in the neuromasts of the posterior lateral line (Figure 2H). The neuromasts are sensory epithelial receptor organs that contain hair cells that respond to changes in movement and pressure of the surrounding water (Chitnis et al., 2012; Thomas et al., 2015). The lipid droplets appear at the edge of the organs, suggesting that they are likely not within the hair cells, but may be located in either the support cells, mantle cells (Chitnis et al., 2012; Thomas et al., 2015) or in the neuromast border cells (Seleit et al., 2017). These findings are consistent with the report of lipid droplets adjacent to neuromasts in zebrafish imaged with lattice light sheet PAINT microscopy (Legant et al., 2016). Crossing the *Fus*(*EGFP-plin2*) line to transgenic reporter lines for the different cell-types (Parinov et al., 2004; Steiner et al., 2014; Thomas and Raible, 2019) will allow the cellular location of these organelles to be identified and may provide insight into the possible role the lipid droplets play in neuromast physiology.

### Additional transgenic lines are also available for overexpression of human PLIN2 and PLIN3

While the knock-in lines are superior for imaging plin2 and plin3 in the zebrafish because they are expressed under the control of the endogenous promoter and regulatory elements, we also have a number of additional Tol2-based transgenic lines available which could be useful in specific contexts or for specific purposes. These lines express human PLIN2 or PLIN3 under the control of the zebrafish FABP2, FABP10a or Hsp70l promoters for over-expression in the yolk syncytial layer & intestine, liver or throughout the larvae following heat shock, respectively (Table 1 & Table 1 - figure supplement 1).

**Table 1:** Comparison of available transgenic perilipin lines.

| Transgenic Line | Promoter | Coding sequence | Tissue Expression |
| --- | --- | --- | --- |
| *Fus(EGFP-plin2)* | Integration into the endogenous plin2 locus | zebrafish plin2<br>ENSDART00000175378.2 | Intestine, liver, adipose, neuromasts, rare LDs in YSL |
| *Tg(FABP2: EGFP-PLIN2)* | Zebrafish intestinal fatty acid binding protein (FABP2) | Human perilipin 2<br>ENST00000276914.7 | Yolk syncytial layer, intestine |
| *Tg(FABP10a: EGFP-PLIN2)* | Zebrafish liver fatty acid binding protein (FABP10a) | Human perilipin 2<br>ENST00000276914.7 | Liver |
| *Tg(Hsp70I: EGFP-PLIN2)* | Zebrafish heat shock cognate 70-kd protein, like | Human perilipin 2<br>ENST00000276914.7 | Widespread tissue expression following heat shock<br>Labeled LDs observed in intestine and liver |
| *Fus(plin3-RFP)* | Integration into the endogenous plin3 locus | zebrafish plin3<br>ENSDART00000100473.5 | Intestine, liver, adipose<br>Cytoplasmic in addition to LDs |
| *Tg(FABP2: PLIN3-EGFP)* | zebrafish intestinal fatty acid binding protein (FABP2) | Human perilipin 3<br>ENST00000221957.9 | Yolk syncytial layer, intestine<br>Cytoplasmic in addition to labeled LDs in intestine |
| *Tg(Hsp70I: PLIN3-EGFP)* | Zebrafish heat shock cognate 70-kd protein, like | Human perilipin 3<br>ENST00000221957.9 | Widespread tissue expression following heat shock; often mosaic<br>Cytoplasmic in addition to labeled LDs in intestine and liver |

In summary, the *Fus*(*EGFP-plin2*) and *Fus*(*plin3-RFP*) knock-in zebrafish lines provide the opportunity to study perilipins and lipid droplet biology *in vivo* at the organelle, cell, tissue, organ and whole animal level. These lines exploit the advantages of the zebrafish model and will be important tools to understand how lipid droplet dynamics are affected by different genetic and physiological manipulations. Future studies with these fish may also help us explain the poorly understood genetic association of the PLIN2 locus with a host of highly prevalent metabolic diseases such as fatty liver, insulin resistance and type 2 diabetes, cardiovascular disease and atherosclerosis (Conte et al., 2016).

## Methods and materials

### Zebrafish husbandry and maintenance

All procedures using zebrafish (*Danio rerio*) were approved by the Carnegie Institution Department of Embryology Animal Care and Use Committee (Protocol #139). Zebrafish stocks (AB line) were maintained at 27°C in a circulating aquarium facility with a 14:10 h light:dark cycle. For propagation and stock maintenance, starting at 5.5 dpf, larvae were fed with GEMMA Micro 75 (Skretting) 3x a day until 14 dpf, GEMMA Micro 150 3x a day + Artemia 1x daily from 15-42 dpf and then GEMMA Micro 500 1x daily supplemented once a week with Artemia. The nutritional content of GEMMA Micro is: 59% Protein 59%; 14% Lipids, 0.2% Fiber; 14% Ash; 1.3% Phosphorus; 1.5% Calcium; 0.7% Sodium; 23000 IU/kg Vitamin A; 2800 IU/kg Vitamin D3; 1000 mg/kg Vitamin C; 400 mg/kg Vitamin E. Embryos were obtained by natural spawning and raised in embryo medium at 28.5°C in culture dishes in an incubator with a 14:10 h light:dark cycle. Zebrafish sex is not determined until the juvenile stage, so sex is not a variable in experiments with embryos and larvae.

### High-fat & high-cholesterol diets

To feed 6 dpf larvae a high-fat, high-cholesterol meal, larvae were immersed in a solution of 5% chicken egg yolk liposomes in embryo media for 1-2 h on an orbital shaker at 29°C as described in (Zeituni and Farber, 2016). Where noted, BODIPY (558/568)-C12 (D3835, Thermo Fisher Scientific) or BODIPY FL-C12 (D3822, Thermo Fisher Scientific) were included in the egg yolk solution at 4 μg/ml. Following feeding, larvae were washed in embryo media, and screened for full guts by examining intestinal opacity under a stereomicroscope. Fed larvae were either maintained in embryo media until imaging, fixed immediately for *in situ* hybridization or guts were extracted and frozen for qRT-PCR. A high-cholesterol diet was made by soaking Gemma Micro 75 in a diethyl ether and cholesterol (Sigma-Aldrich C8667) for a final content of 4% w/w cholesterol after ether evaporation (based on (Stoletov et al., 2009)). Larvae were fed with this high-cholesterol diet 3x daily from 5.5 dpf to 15 dpf where noted.

### *In situ* hybridization

Zebrafish embryos were staged according to (Kimmel et al., 1995) and fixed 4% paraformaldehyde in phosphate buffered saline overnight at 4°C, washed twice with MeOH and stored in MeOH at −20°C. To generate riboprobes, 754 base pairs of the *perilipin 2 (plin2; ENSDARG00000042332; ENSDART00000175378.2* transcript) and 900 base pairs of the zgc:77486 (*perilipin 3; plin3*; ENSDARG00000013711, *ENSDART00000100473.5* transcript)(GRCz11) mRNA sequences were amplified from cDNA using the primers noted in Supplementary Table 1 and TA cloned into the dual promoter pCRII-TOPO^®^ (Thermo Fisher Scientific, K207020). Sense and anti-sense probes were synthesized using the DIG RNA labeling kit (Roche 11277073910) using T7 and SP6 polymerases (Roche 10881767001 & 10810274001). Whole mount in situ hybridization was performed as previously described (Thisse and Thisse, 2008) on 6 dpf unfed and high-fat fed larvae. Larvae were mounted in glycerol and imaged using a Nikon SMZ1500 microscope with HR Plan Apo 1x WD 54 objective, Infinity 3 Lumenera camera and Infinity Analyze 6.5 software or a Nikon E800 microscope with a 20X/0.75 Plan Apo Nikon objective and Canon EOS T3 camera using EOS Utility image acquisition software.

### DNA extraction and genotyping

Genomic DNA was extracted from embryos, larvae or adult fin clips using a modified version of the HotSHOT DNA extraction protocol (Meeker et al., 2007). Embryos or tissue were heated to 95°C for 18 minutes in 100 μL of 50 mM NaOH, cooled to 25°C and neutralized with 10 μL of 1 M Tris-HCL pH 8.0. The gDNA extractions and PCR verifying integration of the fluorescent tags into the genomic loci was performed using the REDExtract-N-Amp Tissue PCR kit (Sigma-Aldrich). PCR amplicons were run on 1 or 2% agarose gels in TBE and gels were imaged with Bio-Rad Gel ChemiDoc XRS system and Quantity One software.

### RNA isolation, cDNA synthesis and quantitative RT-PCR

Following a 90 min feed with 5% chicken egg yolk, guts were dissected from larvae (6 dpf, 10 guts pooled per sample) and stored in RNA*later* (Thermo Fisher Scientific AM7020) at 4°C for 1 week. RNA was isolated using a Trizol-based RNA prep adapted from (N.J. and T.L., 2014). Samples were subsequently treated with DNase I and purified using the RNA Clean and Concentrator kit (Zymo Research R1013). cDNA was synthesized using the iScript™ cDNA Synthesis Kit (1708891, Bio-Rad Laboratories, Inc.). qRT-PCR samples were prepared using SsoAdvanced™ Universal SYBR^®^ Green Supermix (1725271, Bio-Rad Laboratories, Inc.). Primers targeting zebrafish *plin2* transcripts were previously validated (See Supplementary Table 1)(Zeituni et al., 2016) and zebrafish 18S (*rps18*) was used as the reference gene (Otis et al., 2015). qRT-PCR was performed in triplicate for each sample with the Bio-Rad CFX96 Real-Time System with 45 cycles: 95°C for 15 seconds, 59°C for 20 seconds, and 72°C for 20 seconds. Results were analyzed with the Bio-Rad CFX Manager 3.0 software and relative gene expression was calculated using the ΔΔCT method (Livak and Schmittgen, 2001).

### Genome editing to create perilipin fusion lines

*Fus*(*EGFP-plin2*) and *Fus*(*plin3-RFP*) lines were created with TALEN-mediated genome editing. The genomic region around the location targeted for editing in the *plin2* and *plin3* genes was amplified by PCR and sequenced from multiple wild-type AB fish to identify any discrepancies between the published sequences and Farber lab stocks. During this process we identified a variable 54-bp region prior to exon 1 in the *plin2-203* transcript (*ENSDART00000175378.2*)(See Figure 1 – figure supplement 1) and discovered a polymorphism (T>C) in the ATG designated as the start codon in the *plin2-202 ENSDART00000129407.4* transcript. We designed our editing strategy to fuse the EGFP coding sequence in-frame with the *ENSDART00000175378.2* transcript in fish carrying the 54-bp intronic sequence and performed editing only in fish carrying this full-length sequence. Two pairs of TALENs were designed per gene using the Mojo Hand design tool (Neff et al., 2013) and cloned with the FusX assembly system and the pKT3Ts-goldyTALEN vector (Ma et al., 2013; Ma et al., 2016; Welker et al., 2016). The designed TALEN pairs for *plin2* (pair 1 TTTCTGCTAACATGG & AAATAACCAGGTTTGCC; pair 2 TTTCTGCTAACATGGGT & AATAACCAGGTTTGCC) flank a Fok1 restriction site just downstream of the endogenous start codon. The designed TALEN pairs for *plin3* (pair 1 TTGCGCCTCAGATAAC & AATTGCCACACAACCT; pair 2 CAGATAACAGAGAAA & CACACAACCTAAATA) flank a Hph1 site immediately upstream of the endogenous termination codon. TALEN mRNA was in vitro transcribed using the T3 Message Machine Kit (Thermo Fisher Scientific, AM1348), injected into 1-cell stage zebrafish embryos, and cutting efficiency of each pair was assessed by monitoring the loss of either FokI (NEB R0109) or HphI (NEB R0158) digestion due to TALEN nuclease activity. Nuclease activity was higher for *plin2* TALEN pair 1 and *plin3* TALEN pair 1 and these were used subsequently for genome integration. Donor plasmids used as templates for homology directed repair were assembled using the three-fragment MultiSite gateway assembly system (Invitrogen, 12537-023). For *plin2*, the 5′ element consisted of 594bp of genomic sequence upstream of the *plin2* start codon, the middle-entry element contained the *plin2* kozak sequence followed by the EGFP coding sequence lacking a termination codon that was in-frame with the 3′ element which consisted of 900bp of genomic sequence including and downstream of *plin2* start codon. For *plin3*, the 5′ element consisted of the 679 bp of genomic sequence immediately upstream of the termination codon, a middle entry element of in-frame tagRFP-t (amplified from Addgene # 61390, which has been codon modified for zebrafish (Horstick et al., 2015)) with a C-terminal termination codon, and a 3′ element consisting of the 444 bp genomic sequence downstream of the *plin3* termination codon. Genome integration was accomplished by co-injection of 150 pg of TALEN mRNA and 100 pg of donor plasmid into 1-cell stage embryos. Injected embryos were raised to adulthood, out-crossed to wild-type fish and resulting F1 progeny were screened for either EGFP or RFP fluorescence; it was necessary to feed the *Fus*(*EGFP-plin2*) fish with a high-fat meal in order to detect EGFP-plin2 fluorescence when integrated correctly. Correct in-frame integration of the fluorescent reporters was confirmed by PCR and sequencing. *Fus*(*EGFP-plin2*) fish can also be genotyped using primers for EGFP. For primer information, see Supplementary Table 1.

### Additional transgenic zebrafish

Additional transgenic zebrafish expressing human *perilipin 2 (PLIN2, ENSG00000147872 GRCh38.p13*) or *perilipin 3* (*PLIN3, ENSG00000105355*) under the control of various promoters were generated with the Tol2-Gateway molecular cloning system (Kwan et al., 2007). The coding sequence of human *PLIN2* with an N-terminal EGFP tag was provided by John McLauchlan (Targett-Adams et al., 2003) and re-cloned into pCR8 (ThermoFisher Scientific). The human *PLIN3* (TIP47) coding sequence was obtained from Flexgene clones collection (Harvard Medical School, clone ID: HsCD00004695) and re-cloned into pCR8. The intestine-specific intestinal fatty acid binding protein (*fapb2*) promoter was provided by Michel Bagnat; the liver-specific liver fatty acid binding protein 10a (p5E *fabp10a*, (−2.8 kb)), originally described in (Her et al., 2003), was provided by Brian Link. The heat shock cognate 70-kDa protein, like (*hsp70l*) promoter (p5E-hsp70l), p3E-EGFPpA, pME-EGFP no stop, and p3E-polyA plasmids were originally provided by Chi-bin Chien (Kwan et al., 2007). Gateway recombination was used to combine entry plasmids into the pDestTol2Pa2 plasmid to create *tg(fabp2: EGFP-PLIN2), tg(fabp10a: EGFP-PLIN2), tg(hsp70l: EGFP-PLIN2), tg(fabp2: PLIN3-EGFP)* and *tg(hsp70l: PLIN3-EGFP)* transgene constructs. Plasmids were injected (25-50 pg) along with 40 pg tol2 transposase mRNA into 1-cell stage AB embryos. Zebrafish were raised to adulthood, out-crossed to wild-type fish and resulting embryos were screened for progeny stably expressing the fluorescent constructs. Transgenic larvae expressing a heat shock-inducible construct were incubated at 37°C for 45 min in 15 mL of embryo media and screened a few hours later. Embryos expressing a *fabp2*-driven construct were screened at 2-4 dpf for EGFP expression in the yolk syncytial layer and embryos expressing the *fabp10a*-driven construct were screened at 5 or 6-dpf following liver development. At least two stable lines per construct were initially generated, the pattern of expression was verified to be the same in each line and subsequently, a single line for each construct was used for experiments and propagated by out-crossing to wild-type AB fish.

### Fluorescence microscopy

Zebrafish larvae at 6 dpf were mounted in 3% methylcellulose in embryo media on glass slides and imaged live with a Zeiss Axio Zoom V16 microscope equipped with a Zeiss PlanNeoFluar Z 1x/0.25 FWD 56 mm objective, AxioCam MRm camera, EGFP and Cy3 filters and Zen 2.5 Blue edition software. For confocal imaging of lipid droplets in the tissues of live larvae at 6, 7 and 15 dpf, larvae were anesthetized with tricaine (Sigma-Aldrich A5040) and mounted in 3% methylcellulose between on glass slides with bridged coverslips. Images were obtained with a Leica DMI6000 inverted microscope and Leica 63x/1.4 HCX PL Apo oil-immersion objective with a Leica TCS-SP5 II confocal scanner with photomultiplier detectors using Leica Application Suite Advanced Fluorescence 2.7.3.9723 image acquisition software. Images were obtained using 4-line average, and recorded with 12-bit dynamic range. EGFP and BODIPY-FL were excited with an argon laser (488 nm) and had a collection window of 498 – 530 nm. BODIPY (558/568)-C12 was imaged with 561 laser and collection window of 571 – 610 nm and mTagRFP-t was imaged with 561 laser and collection window of 575 – 650 nm.

### Additional software

Graphing was performed with GraphPad Prism (GraphPad Software). DNA, mRNA and protein sequence alignments were performed with MacVector V15.5 (MacVector, Inc.). Micrographs were adjusted and cropped as needed in Fiji (NIH) and figures were assembled in Adobe Illustrator CS5 (Adobe Systems). Microsoft Word and Excel were used for manuscript preparation and data analysis, and references were compiled with EndNote 8x.

## Acknowledgments

We gratefully acknowledge Andrew Rock, Carmen Tull, Julia Baer, and Mackenzie Klemek and Hannah Kozan for fish husbandry, Amy Kowalski, Lamia Wahba, Blake Caldwell, James Thierer and Erin Zeituni for synthesis of various plasmids, Cassandra Bullard & Camden Daby for assembly of TALEN plasmids, Stephanie Yan for technical assistance and David Raible for input regarding neuromasts.

## Author Contributions

M.H.W. contributed to study conception and design, resources, data acquisition, analysis and interpretation of data, data presentation, manuscript writing and revision and funding acquisition. S.C.E. contributed methodology, resources and funding acquisition. S.A.F. contributed to study conception and design, resources, manuscript review and editing, supervision, project administration and funding acquisition.

## Supplementary Figures

**Figure 1 – figure supplement 1:**
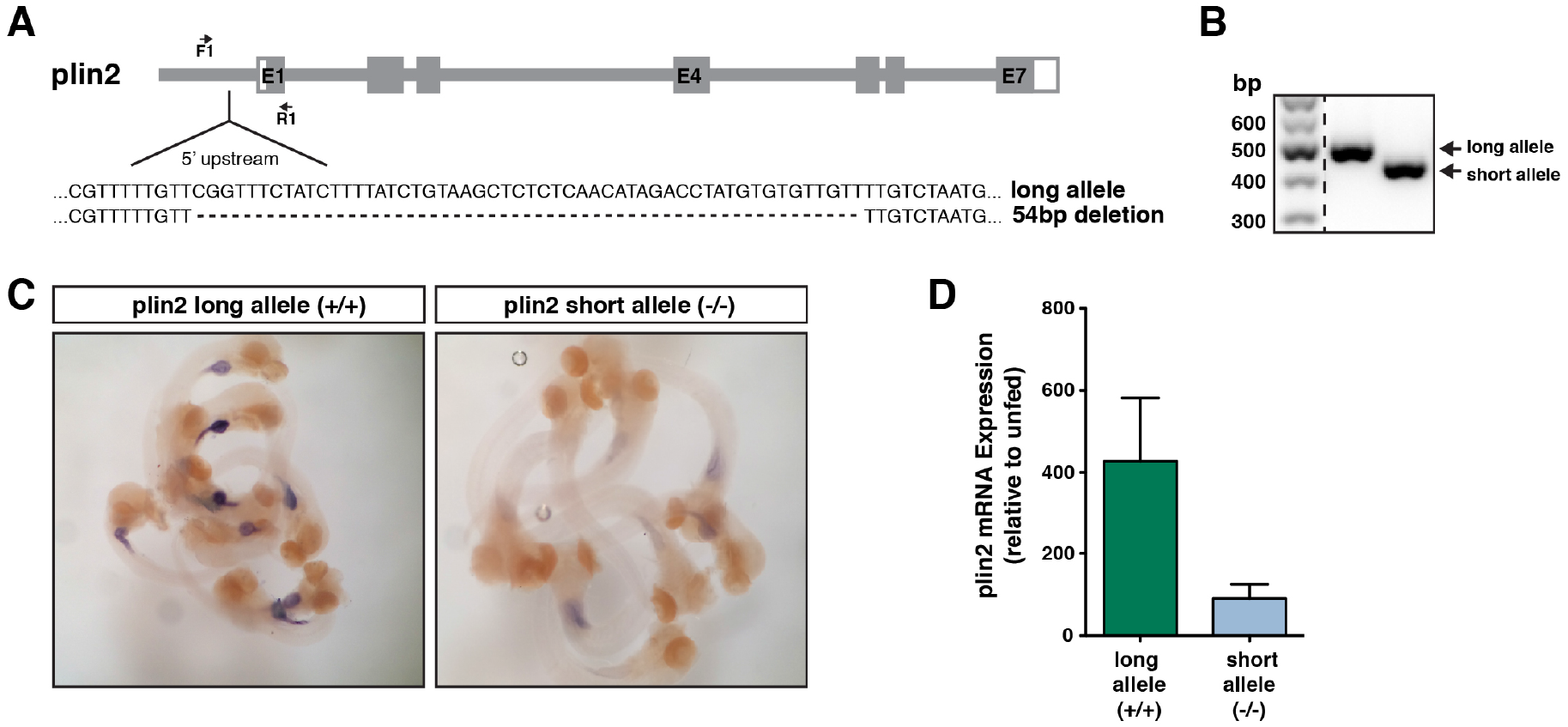
A deletion upstream of exon 1 in plin2 impacts gene expression following a high fat meal. (A) A 54-bp deletion was noted in the 5′ untranslated region upstream of exon 1 in the *plin2 ENSDART00000175378.2* transcript in AB wild-type stocks. (B) RT-PCR using the primers noted in A reveal the long vs. short (deletion) alleles. (C) *In situ* hybridization indicates that larvae homozygous for the long allele have stronger expression of *plin2* in the intestine following a high-fat meal than fish homozygous for the short allele (~10 larvae shown at 6 dpf, fed 90 min prior to fixation). (D) Quantitative RT-PCR confirmed the difference in *plin2* expression induction between larvae with the long vs. short allele following a high fat meal relative to unfed controls (N = 5 samples of isolated guts from 10 larvae per sample following a 90 min feed, samples are from two independent experiments; mean +/- SD).

**Figure 1 – figure supplement 2:**
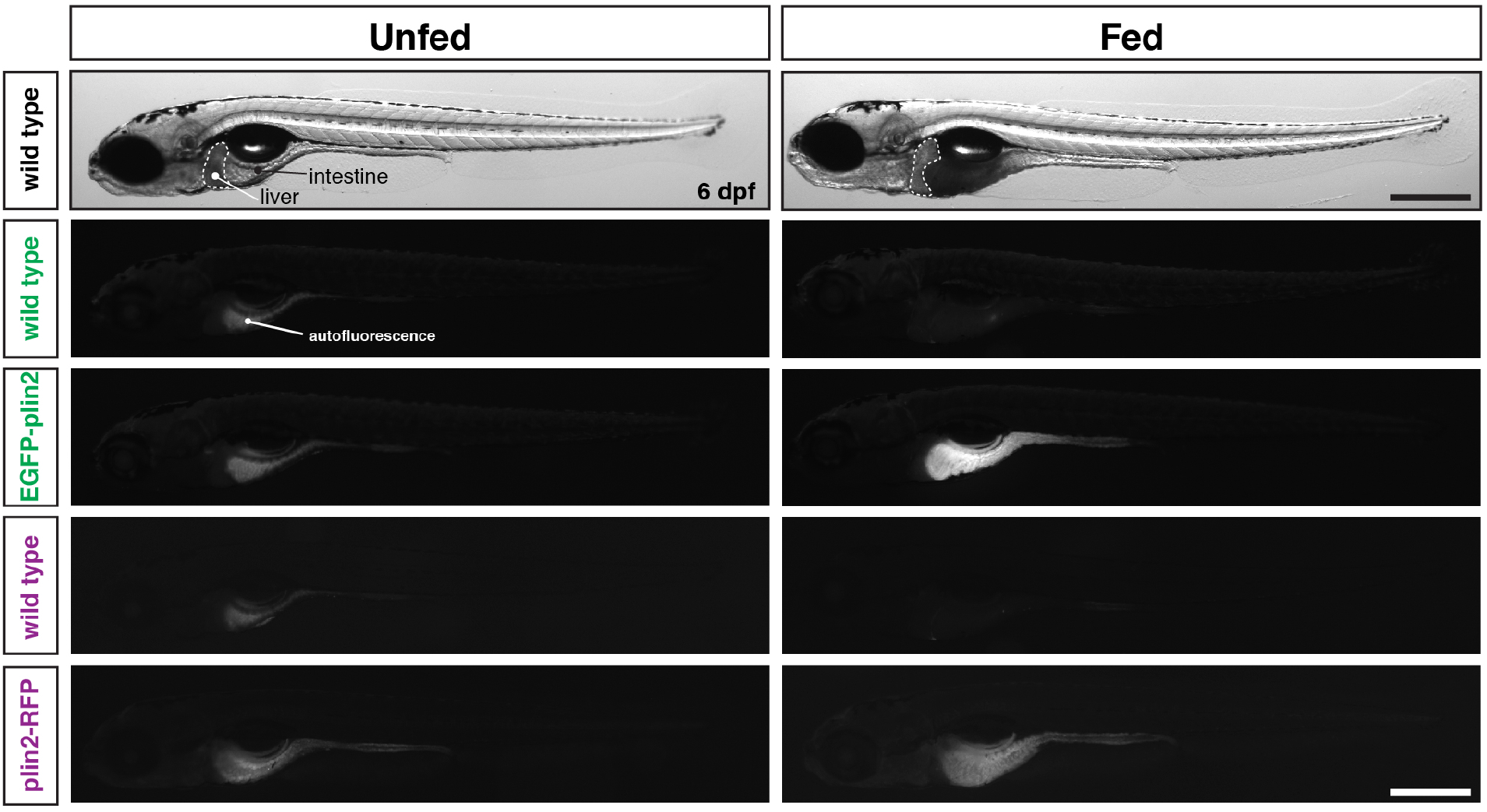
Whole fish images corresponding to Figure 1E. Whole mount images of the same fish shown in figure 1E. Scale = 500 μm.

**Table 1 – figure supplement 1:**
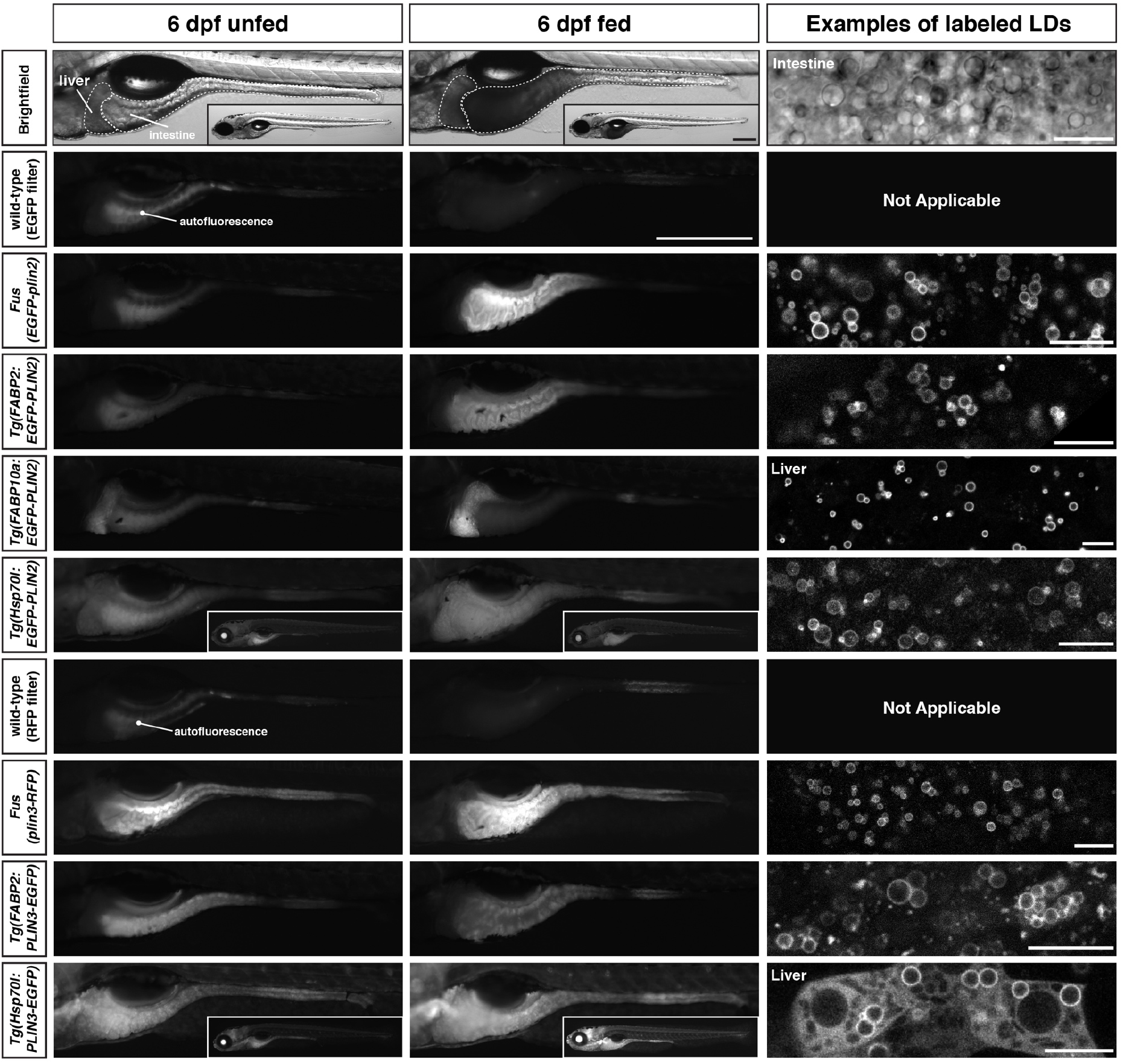
Whole mount images and examples of perilipin-labeled lipid droplets corresponding to the transgenic zebrafish lines noted in Table 1. All fish are heterozygous for the noted transgene. Heat shock transgenic lines were incubated at 37°C for 45 min prior to feeding. For whole mount images, larvae were fed for 2 h with a high-fat meal and imaged 3-4.5 h (plin3 lines) or 5-8 h (plin2 lines) following the start of the feed. Where appropriate, images of whole fish are included as insets. Scale = 500 μm for main images and insets. In the right column, examples of confocal micrographs are included to show the fluorescent perilipin proteins labeling lipid droplets in the various transgenic lines following a high-fat meal. Unless noted, images are from the intestine. Scale = 10 μm for each image.

**Supplementary Table 1:**
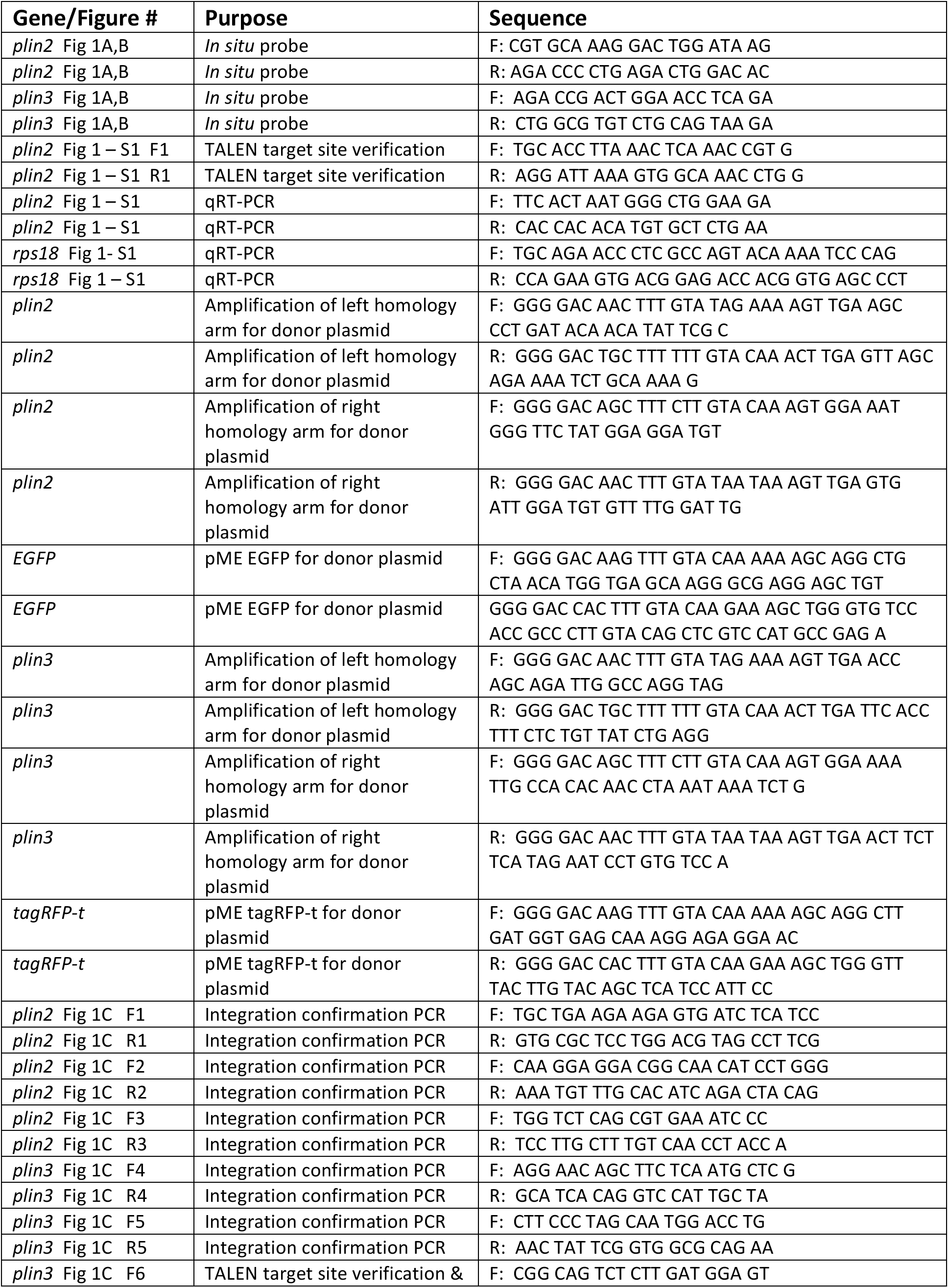
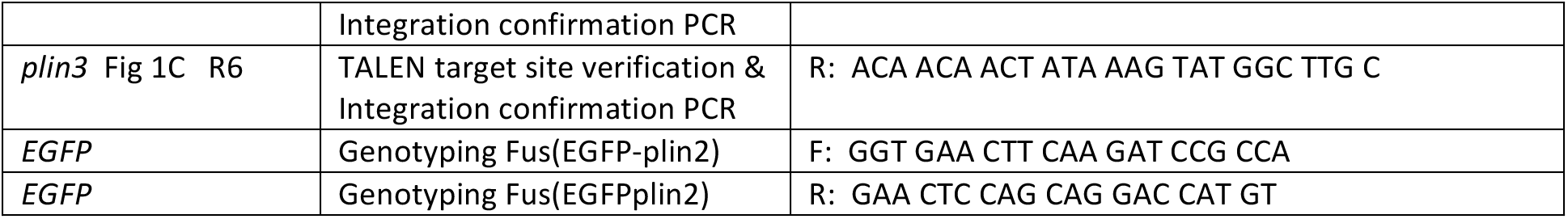
Primers

